# A non-functional copy of the salmonid sex determining gene (*sdY*) is responsible for the “apparent” XY females in Chinook salmon, *Oncorhynchus tshawytscha*

**DOI:** 10.1101/2021.07.28.454148

**Authors:** Sylvain Bertho, Amaury Herpin, Elodie Jouanno, Ayaka Yano, Julien Bobe, Hugues Parrinello, Laurent Journot, René Guyomard, Thomas Muller, Penny Swanson, Garrett McKinney, Kevin Williamson, Mariah Meek, Manfred Schartl, Yann Guiguen

**Affiliations:** INRAE, LPGP, 35000 Rennes, France; Physiological Chemistry, Biocenter, University of Wuerzburg, 97074 Wuerzburg, Germany; Institut de Génomique Fonctionnelle, IGF, CNRS, INSERM, Univ. Montpellier, F-34094 Montpellier, France; GABI, INRAE, AgroParisTech, Université Paris-Saclay, Jouy-en-Josas, France; Julius-von-Sachs-Institute, Department of Molecular Plant Physiology and Biophysics, University of Wuerzburg, D-97082 Wuerzburg, Germany; Environmental and Fisheries Sciences Division, Northwest Fisheries Science Center, National Marine Fisheries Service, National Oceanic and Atmospheric Administration, 2725 Montlake Blvd East, Seattle, WA 98112, USA; Molecular Genetics Laboratory, Washington Department of Fish & Wildlife, 600 Capitol Way N. Olympia, WA 98501; Atreca, Inc., 835 Industrial Road, Suite 400, San Carlos, CA 94070; Michigan State University, Dept. of Integrative Biology, AgBio Research, and Ecology, Evolution, and Behavior Program, 288 Farm Lane, East Lansing, MI 48824; The Xiphophorus Genetic Stock Center, Department of Chemistry and Biochemistry, Texas State University, San Marcos, Texas, USA; Developmental Biochemistry, Biocenter, University of Wuerzburg, 97074 Wuerzburg, Germany

## Abstract

Many salmonids have a male heterogametic (XX/XY) sex determination system, and they are supposed to have a conserved master sex determining gene (*sdY*), that interacts at the protein level with Foxl2 leading to the blockage of the synergistic induction of Foxl2 and Nr5a1 of the *cyp19a1a* promoter. However, this hypothesis of a conserved master sex determining role of *sdY* in salmonids is still challenged by a few exceptions, one of them being the presence of some naturally occurring “apparent” XY Chinook salmon females. Here we show that XY Chinook salmon females have a *sdY* gene (*sdY-N183*), which has one missense mutation leading to a substitution of a conserved isoleucine to an asparagine (SdY I183N). In contrast, Chinook salmon males have both a non-mutated *sdY-I183* gene and the missense mutation *sdY-N183* gene. The 3D model of SdY-N183 predicts that the I183N hydrophobic to hydrophilic amino acid change leads to a local modification of the β-sandwich structure of SdY. Using *in vitro* cell transfection assays we found that SdY-N183, like SdY-I183, is preferentially localized in the cytoplasm. However, compared to SdY-I183, SdY-N183 is more prone to degradation, its nuclear translocation by Foxl2 is reduced and SdY-N183 is unable to significantly repress the synergistic Foxl2/Nr5a1 induction of the *cyp19a1a* promoter. Altogether our results suggest that the *sdY-N183* gene of XY Chinook females is a non-functional gene and that SdY-N183 is no longer able to promote testicular differentiation by impairing the synthesis of estrogens in the early differentiating gonads of wild Chinook salmon XY females.

## INTRODUCTION

Genetic sex determination is a widespread mechanism in vertebrates controlled by master sex determining genes acting on the top of a genetic cascade, ultimately leading to male and female phenotypes (Bachtrog *et al*. 2014). Despite recent technological improvements in genome sequencing and genetics, and despite increasing discoveries of master sex-determining genes or candidates in vertebrates only a handful have been functionally characterized to a certain extent. In fish most of the currently known sex determining genes are poorly conserved; like for instance *dmrt1bY* that is only found in *Oryzias latipes* and *O. curvinotus* (Schartl 2004) or *amhr2Y* in some *Takifugu* species (Ieda *et al*. 2018). In contrast, most salmonids have been found to harbor the same unusual sex determining gene named *sdY* (*sexually dimorphic on the* Y). In rainbow trout (*Oncorhynchus mykiss*) this gene, which arose from a duplication of the *irf9* immune related gene, is necessary and sufficient to drive testicular differentiation (Yano *et al*. 2012, 2014). SdY triggers its action by interacting with the conserved female differentiation factor Foxl2 (Bertho *et al*. 2016), ultimately preventing the regulation of estrogen synthesis needed for ovarian differentiation (Bertho *et al*. 2018). As *sdY* is genotypically tightly sex-linked to male development in most salmonid species, it has been suggested that *sdY* could have been conserved over 50-90 million years as the only sex determining gene of all extant salmonids (Yano *et al*. 2013). However, this evolutionary conservation hypothesis has been challenged by some unresolved exceptions to the rule (Yano *et al*. 2013; Cavileer *et al*. 2015; Larson *et al*. 2016; Podlesnykh *et al*. 2017; Ayllon *et al*. 2020; Brown *et al*. 2020) suggesting that *sdY* can be non-functional in some salmonids, or that environmental factors override the function of *sdY*. In cases of *sdY* negative males, the sex-linkage discrepancies could be explained to be the result from neo-masculinization of XX females. This phenomenon has been reported in many fish species including some salmonids (Quillet *et al*. 2002; Valdivia *et al*. 2013). But some studies also report the existence of *sdY* positive females. This is more difficult to reconcile with the idea that *sdY* is still acting as a male sex determining gene in these species. One of the best documented case of such exceptions to the rule is the existence of “apparent” XY females in wild populations of Chinook salmon, *Oncorhynchus tshawytscha*. In this species discrepancies between genotypic and phenotypic sex have been found with some phenotypic females being described with a male genotype, as deduced from the presence of the male specific marker, *OtY1* (Williamson and May 2002). These XY females are fully fertile and cannot be distinguished phenotypically from genetically normal XX females (Williamson and May, 2002). This observation has been reported several times and in different Northwest Pacific regions including the Columbia river (Nagler *et al*. 2001; Chowen and Nagler 2004), Alaska (Yano *et al*. 2013; Cavileer *et al*. 2015), Idaho, Washington (Cavileer *et al*. 2015), and California (Williamson and May 2002, 2005; Williamson *et al*. 2008). The incidence of these XY females varies between 20 and 38% in Central Valley rivers while ranging between 0 and 14% under hatchery conditions (Williamson and May 2002). In addition, independent surveys found proportions of wild-caught XY females ranging from 12% (Cavileer *et al*. 2015) up to 84% (Nagler *et al*. 2001). These studies indeed raised many important concerns about the underlying mechanisms of the observed “outliers” and their impact on wild and hatchery Chinook salmon populations. Multiple, independent hypotheses were proposed to explain this genotype/phenotype incongruence, including the possibility that Chinook salmon could be feminized due to endocrine-disruptor chemicals (EDCs) or pollutant exposition (Nagler *et al*. 2001). Such hypotheses were later excluded using artificial crosses between genotypically normal males (XY) and XY females, showing that half of their phenotypic female offspring were also XY females (Williamson and May 2005) based on Y-chromosome markers (Du *et al*. 1993; Noakes and Phillips 2003). In addition, fluorescence *in situ* hybridization revealed that XY-female Chinook salmon in California are not the product of a Y chromosome to autosome translocation (Williamson *et al*. 2008) and that these XY females are positive for the *sdY* gene (Cavileer *et al*. 2015).

We explored sex determination in these “apparent” XY Chinook salmon females to investigate if *sdY* could be still considered as the master sex determining gene in this species despite the existence of *sdY* positive phenotypic females. We amplified and sequenced *sdY* gene between the exon 2 and 3 of XY Chinook salmon females and found they have a missense mutation in the third exon of the *sdY* gene that produces a single amino acid change (I183N) in a highly conserved position of the SdY protein, while males have the wildtype copy of SdY. The mutation modifies the three-dimensional (3D) structure. The SdY-N183 protein is less stable than the wild-type SdY (SdY-I183) and is affected in its ability to interact with its protein partner, Foxl2 (Bertho *et al*. 2018). This failure in turn leads to the inability to repress the *cyp19a1a* promoter and thereby to suppress female development. Altogether our results suggest that the *sdY-N183* copy in XY Chinook salmon females is inactive and cannot block the female pathway in the same way as the wildtype *sdY* gene. Our results provide an explanation for the existence of naturally occurring XY Chinook salmon females and support the role of *sdY* as the master male sex determining gene of Chinook salmon.

## RESULTS

### XY Chinook salmon females bear a missense mutated copy of *sdY*

The salmonid male sex determining gene, *sdY*, being only present on the Y chromosome is generally a single copy gene (Yano *et al*. 2012, 2013). Whole transcriptome sequencing of a male Chinook salmon testis, however, revealed two single nucleotide variations (SNVs) in the *sdY* coding region (Figure 1A-1B), suggesting the existence of multiple *sdY* genes or *sdY* alleles. The first SNV is a synonymous A to G transition in exon 2 and the second one an A to T transversion in exon 3 that leads to an amino acid change of isoleucine (I) to asparagine (N). To better understand the relation between sex-phenotypes and *sdY* genotypes in Chinook salmon, we then checked the presence of *sdY* in XX females, XY females and XY males using samples from previously described selective crosses between XY or normal XX females with normal XY males (Williamson and May 2005). In agreements with results from wild-caught Chinook salmon (Cavileer *et al*. 2015), we found that XY females were always *sdY* positive (Supplementary Table 1). Systematic re-sequencing of XY females revealed that they carried the *sdY* SNVs found in the male testis mRNA in exons 2 and 3 (G/G in exon 2 and T/T in exon 3). In contrast all males had a double peak at these positions, i.e., A/G in exon 2 and A/T in exon 3, indicating presence of both *sdY* versions (Figure 1C-C’). However, this does not seem to be an indication of allelic variation of a single *sdY* gene as both versions are present in the XY male offspring of a cross of a XY sire with a normal *sdY* negative XX dam and are not segregating in a Mendelian way in the male offspring. (Supplementary Table 1). This suggests the existence of two *sdY* genes in XY males that could be tightly linked together on the Y chromosome (Figure 2) as almost no recombination was observed in all males (one homozygote on 81 males) from all the three tested progenies. In summary, two version of *sdY* exist in Chinook salmon. The wildtype version would be present only on the Y of males (Y+), while the mutant version would be duplicated on the Y+ of males and present in a single copy on the “apparent” Y (Y-) of XY females (Figure 2).

**Figure 1.**
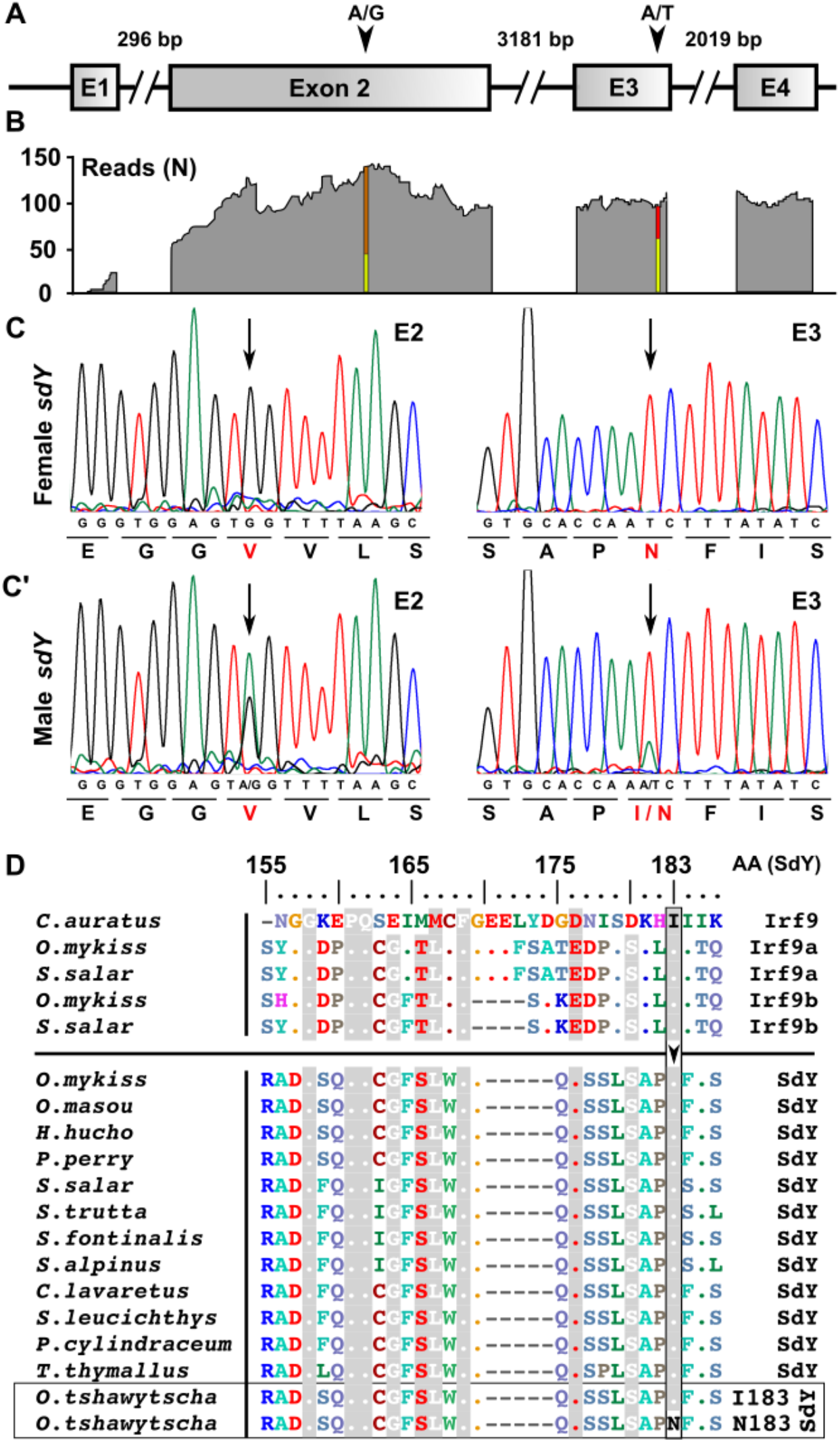
XY Chinook salmon females have a missense mutation in a conserved position of the *sdY* coding sequence. (**A**) Schematic representation of Chinook salmon *SdY* sequence with its 4 exons depicted as square boxes (E1-E4) and the introns as broken lines with intron sizes (bp). (**B**). Remapping of transcriptome reads (N = number of raw remapped reads) from a chinook male testis revealed two SNVs (A/G and A/T) in the coding region of the *sdY* gene. (**C, C’**). Representative sequencing chromatograms of parts of the genomic *sdY* coding sequencing containing SNVs in XY females (**C**) and XY males (**C’**) leading to a synonymous mutation in exon 2 (A/G) and a missense mutation in exon 3 (A/T). (**D**). Alignment of Irf9a, Irf9b and SdY protein sequences in different salmonids species showing the conservation of isoleucine 183 (I) highlighted in grey color and its modification to asparagine (N) only in XY Chinook salmon females (SdY-N183).

**Figure 2.**
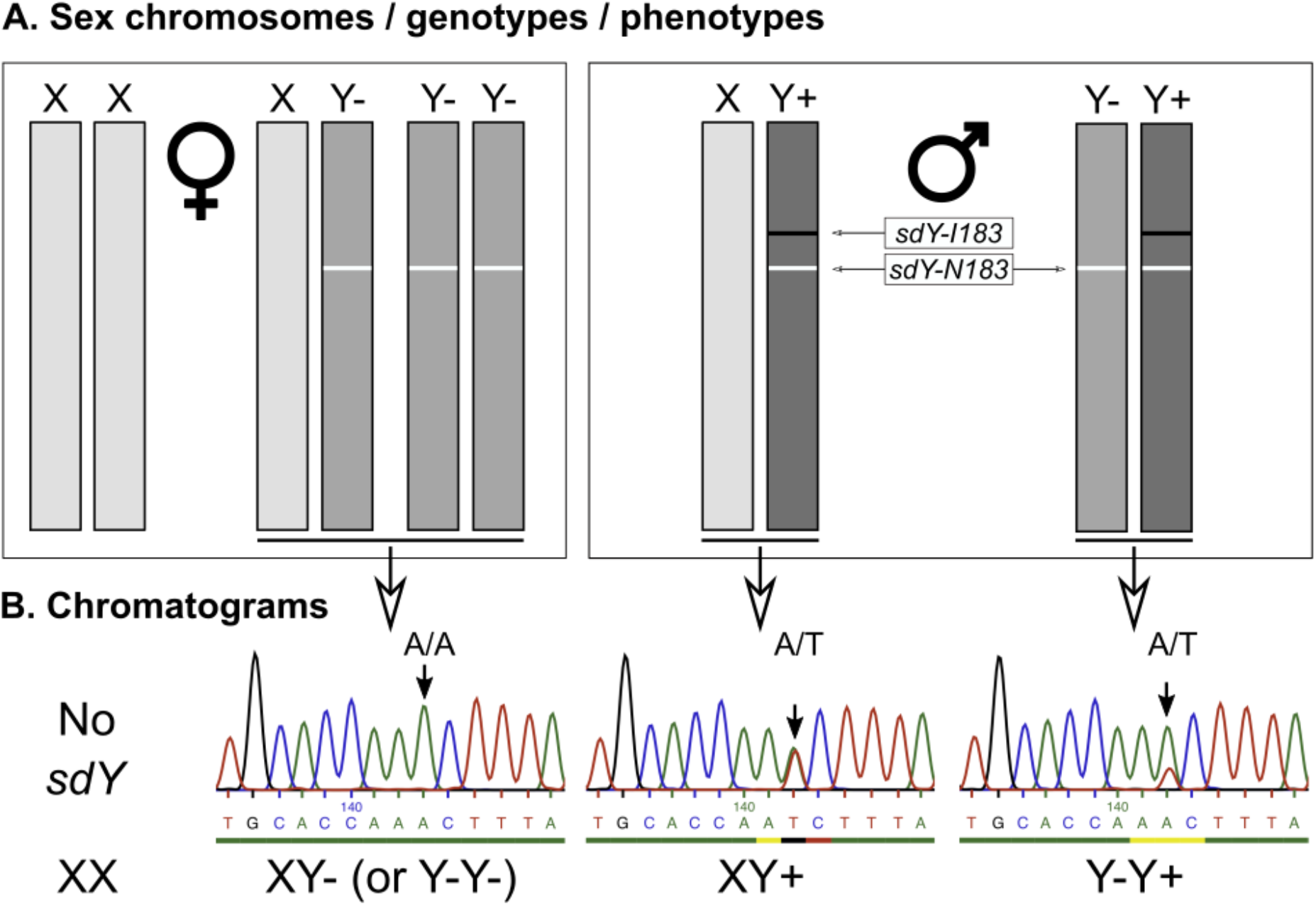
Sex chromosomes, sex genotypes and sex phenotypes in Chinook salmon. (**A**) Schematic representation of sex chromosomes and the hypothetical relation between sex genotypes and sex phenotypes in Chinook salmon. According to our model, phenotypic females can be normal XX females or XY females (XY-) bearing a Y chromosome (Y-) with a single copy *sdY-N183* gene. Phenotypic males can be XY males (XY+) bearing a Y chromosome (Y+) with two copies of the *sdY* gene i.e., *sdY-I183* and *sdY-N183,* or Y-Y+ resulting from the crossing of an XY+ male with an XY- female. In turn, a Y-Y+ males crossed with an XY- female can also generate Y-Y- phenotypic females. (**B**). Representative chromatograms of the sequences around the *sdY* I183N mutation (exon 3) in Chinook salmon. XY females (XY-) are homozygotes A/A for the I183N mutation and males are heterozygotes (A/T). Y-Y- females cannot be discriminated from XY- females based on the chromatogram analysis (single A peak of homozygosity in both cases), but XY+ and Y-Y+ could be in theory identified based on the relative peak height of the A/T “pseudo” alleles. With a 1:1 ratio of *sdY-I183* and *sdY-N183*, XY+ males should have an equal A/T peak height and Y-Y+ with a 1:2 ratio of *sdY-I183* and *sdY-N183* should have an A peak height double from the T peak at the same position. Such chromatogram examples are shown in panel B but due to potential variability of the sequencing reactions this genotyping approach was not retained as an accurate approach to discriminate XY- males from Y-Y+ males in our analyses.

The A to T substitution in exon 3 leads to a transition from an isoleucine (I183) to an asparagine (N183) at amino acid (AA) position 183 of the SdY sequence. The comparison of all SdY protein sequences and some Irf9 protein sequences (Figure 1D) available from salmonids, show that I183 is highly conserved in both SdY and Irf9, suggesting that it could play an important role in SdY function. These results prompted us to explore if the SdY-I183N mutation could be responsible for the phenotype/genotype discrepancy observed in XY females Chinook salmon.

### The I183N substitution predicts potential local SdY misfolding

To examine more precisely what conformational changes are produced by the I183N substitution, we modelled the SdY-N183 protein 3D structure using the IRF5 domain (PDB code 3dsh) as a template (Figure 3A). The model revealed that the I183N substitution is localized at the amino terminal end of the β7-strand shaping the hydrophobic β-sandwich core element of the protein (Figure 3B). The mutation induces a hydrophobic (I) to hydrophilic (N) amino acid pattern change likely modifying at least locally the folding of the β-sandwich. We then tried to find the most suitable model of SdY-N183 in which the hydrophilic side chain may have less negative impact on the folding and protein stability. Being exclusively surrounded by hydrophobic amino acids and with a similar size as I183, it has not been possible to model the N183 in an energetically favorable state. The model also suggests that this unfavorable state might impact the local folding of the β-sandwich and the α_1_-helix. Taken together, our protein structure modelling revealed that the I183N substitution affects the local environment of the β-sandwich potentially disturbing SdY folding and leading to a more unstable protein.

**Figure 3.**
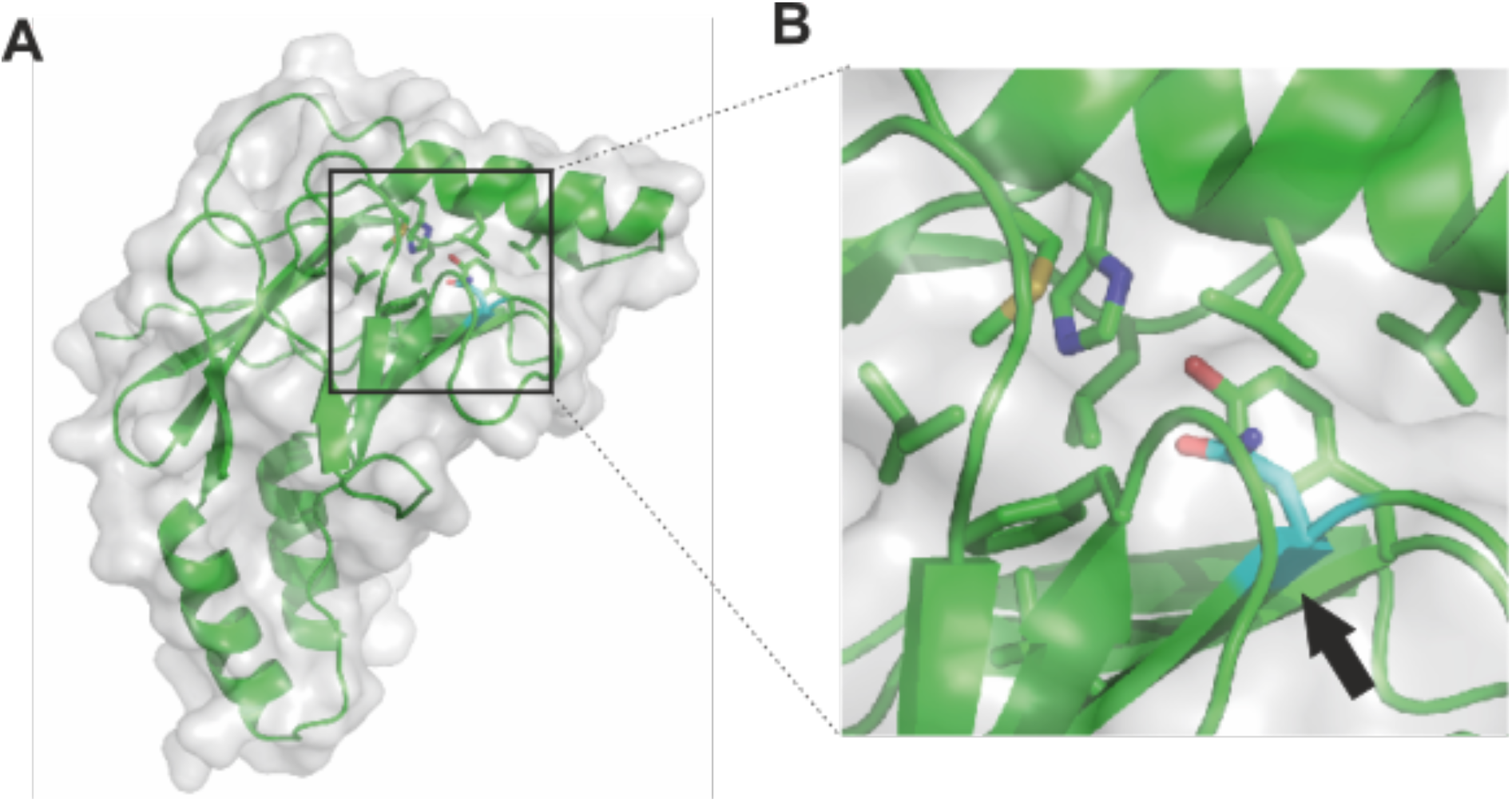
The I183N SdY mutation affects locally the structure of SdY. (**A**) Model of SdY I183N (green) deduced from the protein-protein interaction domain template of IRF5 (PDB ID 3DSH) embedded in the surface representation (grey). (**B**) Magnification around the asparagine residue (N183) in cyan indicated by a black arrow. The mutation is located at the beginning of the β_7_-strand embedded in a hydrophobic pocket leading to a local misfolding.

### SdY-I183N interaction with Foxl2 is reduced compared to wildtype SdY

As Foxl2 has been previously shown to promote SdY translocation from the cytoplasm to the nucleus (Bertho *et al*. 2018), we further investigated the impact of the I183N substitution on the subcellular localization of SdY in presence or absence of Foxl2. For this purpose, we engineered the Chinook salmon mutation into the rainbow trout SdY protein (rtSdY-I183N). Like the wildtype SdY, SdY-I183N was predominantly localized in the cytoplasm when transfected alone into human embryonic kidney (HEK 293T) cells (Figure 4A-4A”, 4E). In contrast to the wildtype protein (Bertho *et al*. 2018), SdY-I183N was also detected in some transfected cells with a nucleo-cytoplasmic localization and even in some cases with a strict nuclear localization (Figure 4B-4B”, 4E). After co-transfection with Foxl2b2, SdY-I183N remained predominantly in the cytoplasmic compartment (Figure 4B-4B”) with a slightly higher percentage of nucleo-cytoplasmic localization (Figure 3C-3C”) compared to transfections of SdY-I183N alone (Figure 4E). Altogether these data show that the cellular localization of SdY-I183N is less restricted than the previously described exclusive cytoplasmic localization of the wildtype protein (Bertho *et al*. 2018), and that SdY-I183N is also strongly impaired in its ability to be translocated into the nucleus by interaction with Foxl2b2 compared to its wildtype counterpart.

**Figure 4.**
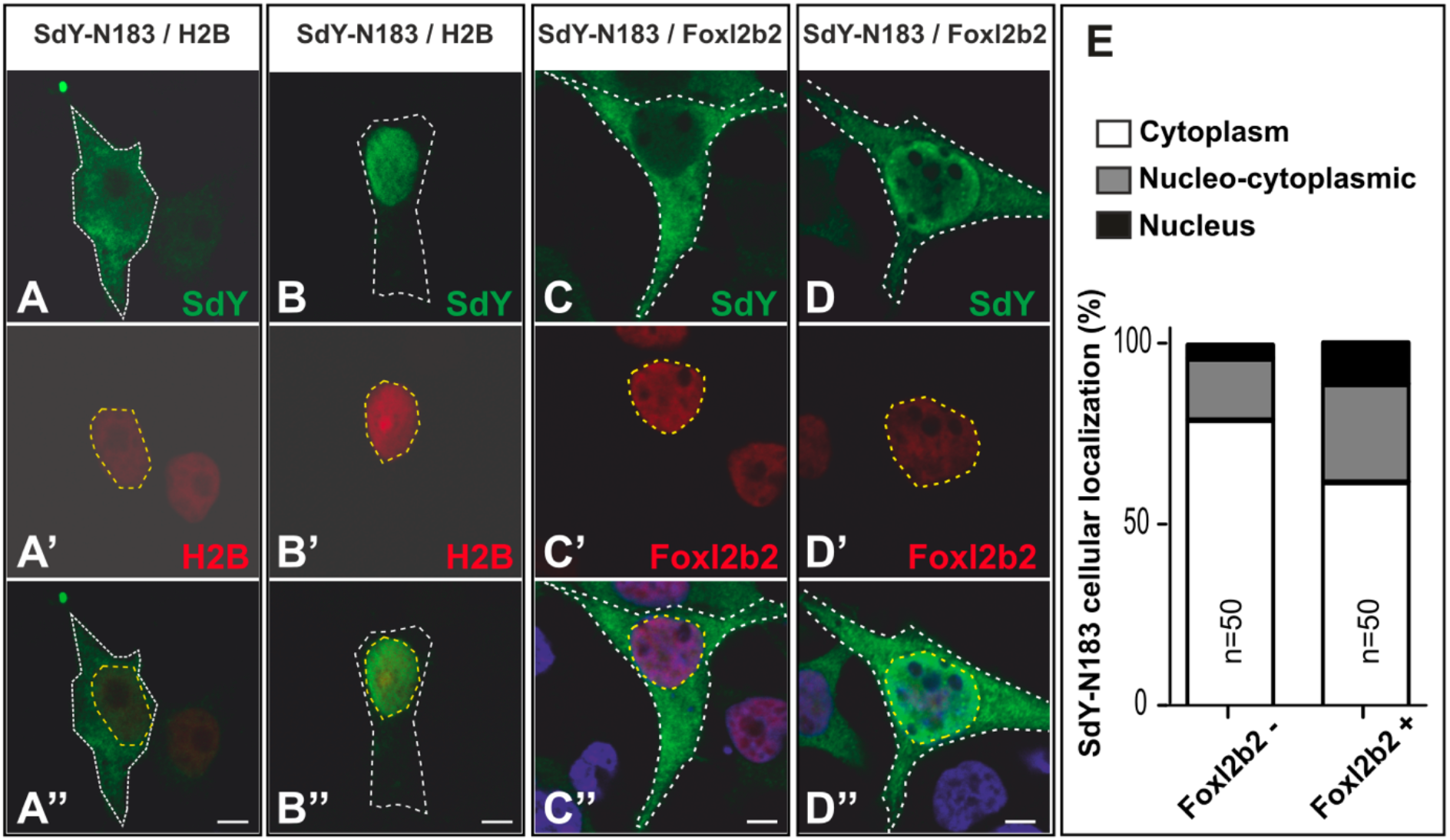
SdY-I183N is localized predominantly in the cytoplasm, and is only slightly translocated in the nucleus after co-transfection with Foxl2b2. SdY-N183 alone is mainly detected in the cytoplasm (**A-A”**) with some transfected cells, however, showing a nucleo-cytoplasmic localization (see panel E for quantification of the different localization percentage) and even in some cells a restricted localization in the nucleus (**B-B”**). After co-transfection with Foxl2b2, SdY-N183 remains also mostly cytoplasmic (**C-C”**) with more transfected cells showing a nucleo-cytoplasmic localization (**D-D”** and panel E for quantification of the different localization percentage) and a complete localization in the nucleus. (**E**) Quantification of the percentage of transfected cells (measured on 50 transfected cells) with a SdY-N183 localization in the cytoplasm (white bar), in the nucleus (black bar) or with a nucleo-cytoplasmic localization (grey bar) with (Foxl2b2 +) or without (Foxl2b2 -) co transfection with Foxl2b2. Human Embryonic Kidney cells (HEK 293T) were transiently co-transfected with rainbow trout SdY-N183 in fusion with 3xFlag tag either with a nucleus marker i.e., Histone H2B-mCherry (H2B), or a rainbow trout Foxl2b2-mCherry expression construct. Rainbow trout SdY-N183 was detected with a FLAG antibody and the nucleus was stain in red for the H2B construct (A’ and B’) or in blue with Hoechst (C’’ and D’’). Scale Bar = 5 μm (A”-D”).

### SdY-I183N is unstable even in presence of Foxl2b2

Because of the local misfolding of SdY-I183N and its lower nuclear translocation following interaction with Foxl2b2, we evaluated the stability of the wildtype and mutant proteins, in presence or absence of Foxl2b2, by time course treatments with a protein synthesis inhibitor (cycloheximide, Figure 5A-5B) and a proteasome inhibitor (MG132, Figure 5C). Both wildtype protein and SdY-I183N expression levels showed a marked decrease 4 hours after the beginning of the cycloheximide treatment (Figure 5A-5A’). Compared to the wildtype protein, SdY-I183N showed reduced expression levels at 4 and 8 hours after cycloheximide treatment (Figure 5A-5A’). After co-transfection with Foxl2b2, expression of both proteins was maintained at relatively high levels 4-hours after the beginning of cycloheximide treatment (Figure 5B-5B’). However, in contrast to the wildtype protein that remained highly expressed at 8 hours post-treatment, SdY-I183N expression dramatically decreased (Figure 5B-5B’). After treatment with the proteasome inhibitor expression levels of both proteins were roughly doubled (Figure 5C-5C’). In the presence of Foxl2b2, wildtype SdY protein expression levels were not increased by the proteasome inhibitor treatment. In contrast, SdY-I183N protein expression level was increased 6-fold relative to untreated cells by the treatment (Figure 5C-5C’). Collectively, this shows that the wildtype SdY protein is stabilized in presence of Foxl2b2, because most likely the interaction with Foxl2b2 protects it from proteasome-mediated degradation. In contrast, the SdY-I183N protein is much more instable probably because of reduced interaction with Foxl2b2 leading to a higher proteasomal degradation.

**Figure 5.**
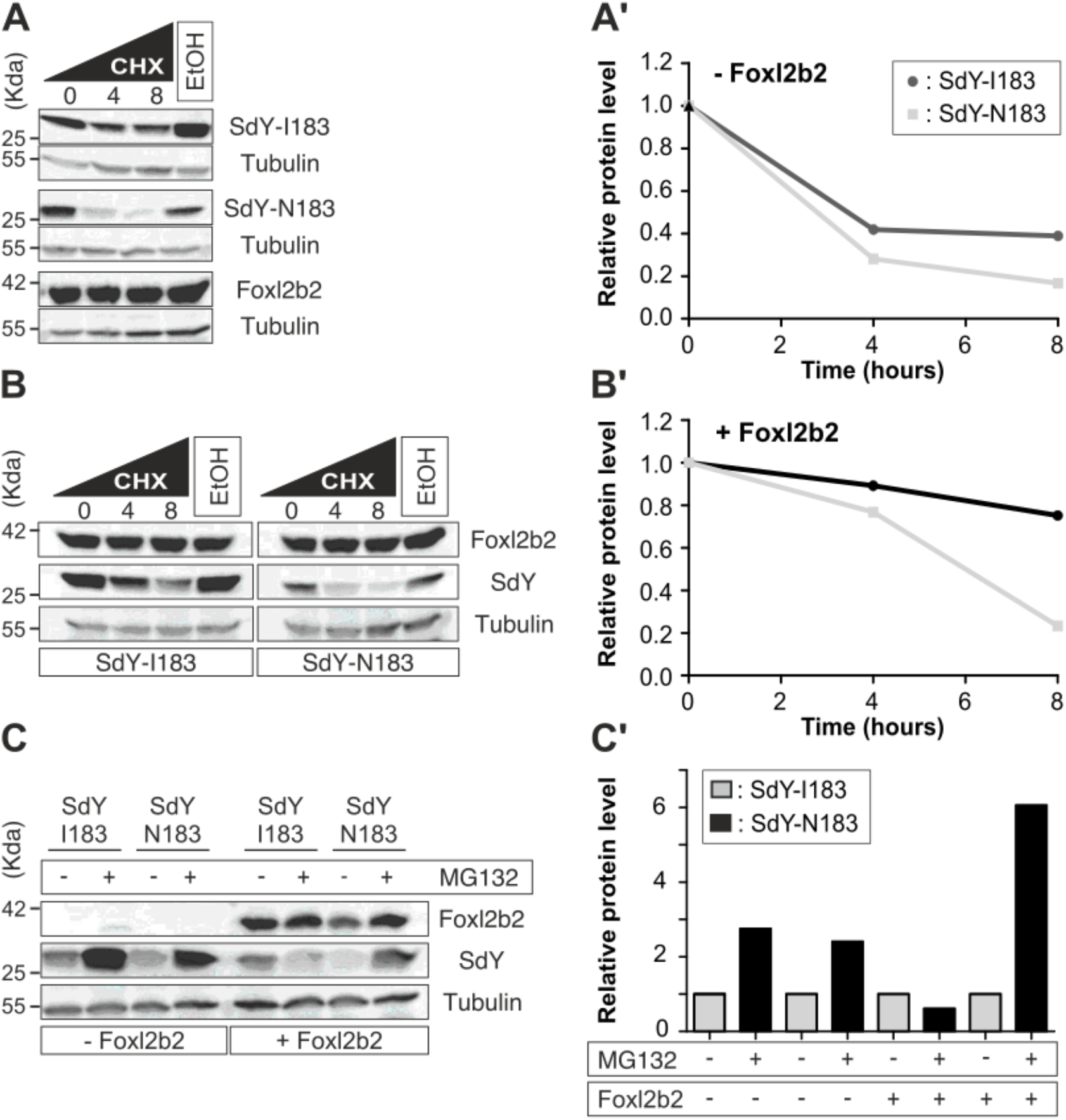
SdY-I183N is unstable even in presence of Foxl2b2. Cycloheximide (CHX) time course were performed to assess SdY-I183 or SdY-N183 stability in presence or absence of Foxl2b2. HEK cells were transiently transfected with SdY-I183, SdY-N183 or Foxl2b2 (**A-A’**) alone or with SdY-I183 or SdY-N183 in combination with Foxl2b2 (**B-B’**). Cells were treated with 50 μm of CHX and harvest at 4 and 8 h (**A’** and **B’**). Lysates were standardized for total protein concentration and expression levels of SdY-I183, SdY-N183 or Foxl2b2 were detected by Western blotting. Tubulin was blotted as a loading control. Foxl2b2 increased SdY-I183 but not SdY-N183 stability (**B**) and (**D**). (**E**). Western blot analysis of SdY-I183, SdY-N183 alone or in combination with Foxl2b2 protein levels following 8 h treatment with proteasome inhibitor MG132. Cells were treated with DMSO (vehicle (control), indicated by a - sign), MG132 (20 μm, indicated by a + sign). Tubulin was blotted as a loading control. (**F**). Quantification of **(E**).

### SdY-I183N is unable to repress the *cyp19a1a* promoter

To get more insight about how the functionality of SdY-I183N is compromised, we also explored the ability of SdY to repress the *cyp19a1a* promoter in synergy with Foxl2 and Nr5a1 (Bertho *et al*. 2018). Like wildtype SdY, SdY-I183N is not able to repress the *cyp19a1a* promoter either with Foxl2 alone or Nr5a1 alone (Figure 6). However, unlike SdY (Supp. Fig 1), SdY-I183N is unable to repress the Foxl2 and Nr5a1 synergetic activation of the *cyp19a1a* promoter (Figure 6), suggesting that *sdY-I183N* is a non-functional master sex-determining gene.

**Figure 6.**
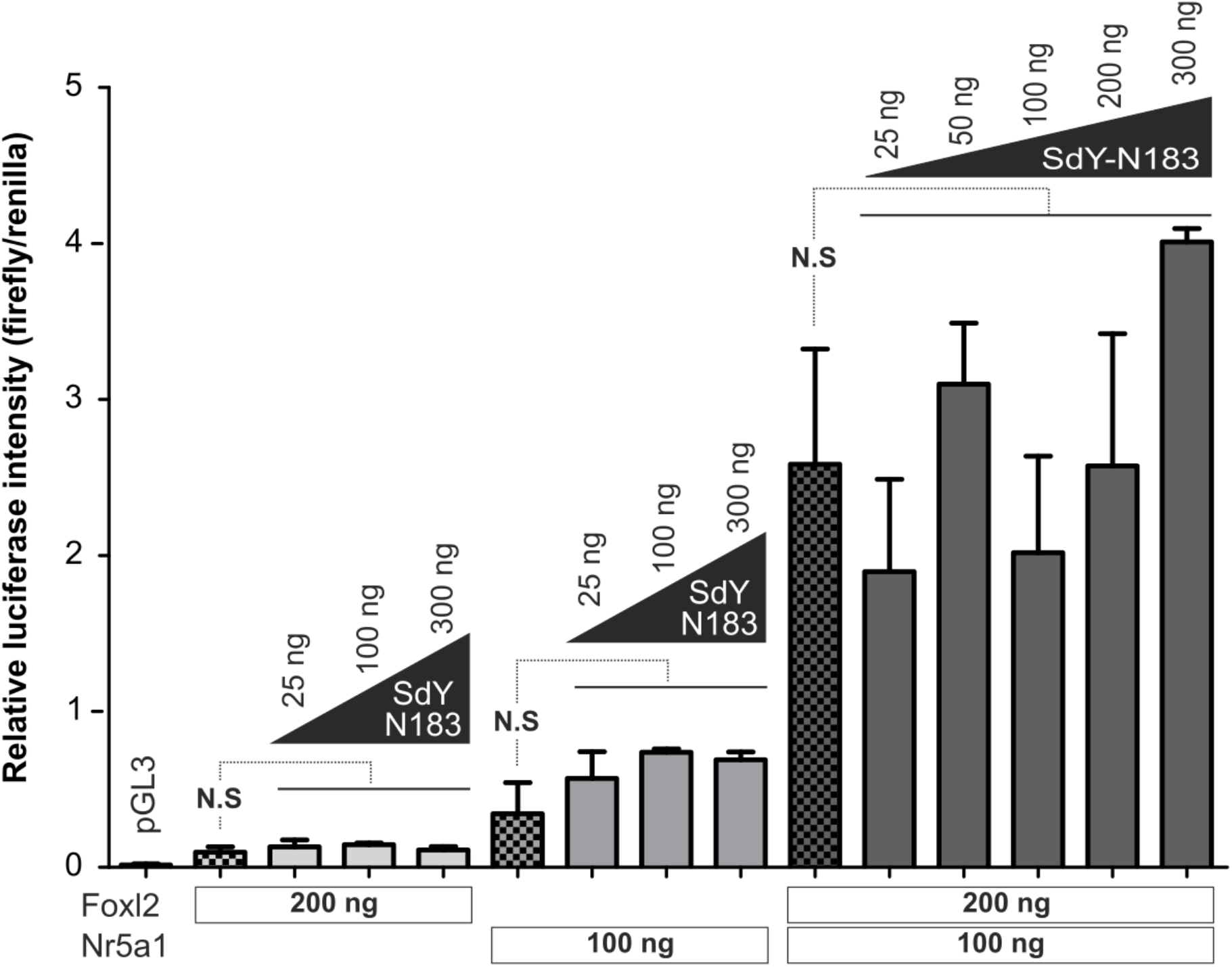
SdY-N183 does not prevent Foxl2/Nr5a1 positive regulation of the *cypl9ala* promoter. (See also supplemental figure 1). The *cyp19a1a* promoter activity *(cyp19a1a* promoter coupled to firefly luciferase) was measured in HEK 293 cells using a luciferase reporter assay and co-transfection of fixed quantities of *nr5a1* (100 ng), *foxl2* (200 ng) and variable quantities (25-300 ng) of *sdY-N183*. Results are calculated from the mean ± SEM of three biological replicates in one experiment. Statistics were calculated with a one-way ANOVA with post-hoc Dunnett tests. N.S: not statistically significant. Empty vector control (pGL3).

## DISCUSSION

Our study identified a missense mutation affecting the sex determining factor SdY in wild XY female Chinook salmon population. SdY-I183N is characterized by a predicted local conformational change in the β-core sandwich, preferential cytoplasmic localization, reduced half-life time, and a lower Foxl2 affinity relative to the wildtype version leading to its inability to repress the *cyp19a1a* promoter.

The isoleucine amino acid at position 183 within the C-terminal domain has a high degree of conservation among the SdY proteins. It is positioned in the protein-protein interaction domain (Interferon–associated domain) of its progenitor Irf9 (Yano et al., 2012; Yano et al., 2013). Previous experiments of genetic ablation of *sdY* using zinc fingers nucleases targeting exon 2 resulted in 14 different mutations such as deletion of leucine 43 (L43) that did not lead to sex reversal while the 13 others mutations lead to a clear male to female sex reversal (Yano et al., 2014). L43 is present in a linker between two β-sheets but not in the β-sandwich. Both amino acids are also conserved in the IAD sequence of Irf9 sequence pointing out a divergence between the conserved primary sequence and the three-dimensional structure. However, this study and our results suggest some crucial amino acid essential for the three-dimensional structure and for the interaction. Such mutations affecting *irf9* have not been described so far.

Taken together, less colocalization with Foxl2 and interaction with lower affinity may be due to the instability of SdY-N183 compared to the wildtype version. Ultimately, the mutated sex determining factor SdY-N183 was not able to act accurately as a repressor of Foxl2 activity. Consistently with these data, SdY-I183N in XY females would not be effective for inducing testicular differentiation.

Here, we bring a potential explanation for the natural sex reversal observed in some wild Chinook salmon populations. Wild fish sex reversals have also been discovered in other species such as Japanese medaka (*O. latipes*) (Matsuda et al., 2002; Otake et al., 2008; Shinomiya et al., 2004). In medaka, two types of mutations affected the sex determining gene *dmy/dmrt1bY* and lead to a XY male to female sex reversal. One type of mutations triggered a low expression of *dmrt1bY* insufficient to tilt the balance toward testis development and the second mutation type affected the amino acid sequence leading to a frameshift and an inactivated Dmrt1bY protein (Otake et al., 2008). Naturally occurring sex reversals were also observed in Nile tilapia (*O. niloticus*) in different Kenyan lakes (Baroiller and D’Cotta, 2016). In pejerrey (*O. bonariensis*) from the Lake Kasumigaura in Japan (Yamamoto et al., 2014) XY female sex reversals were also reported from wild populations. However, the responsible molecular mechanisms have not been revealed yet.

Interestingly, some incongruences between the genotype and phenotype were also described in wild populations in another salmonid, the Sockeye salmon (*O. nerka*), (Larson et al., 2016). In that species, a deeper analysis of the *sdY* sequence should reveal if a mutation is responsible for the observed genotype/phenotype mismatch.

Of note, the SdY-I183N mutation has been evaluated in our study at the individual scale within a few families, but its frequency at the population level has not been thoroughly characterized in Chinook salmon. Further information would then be needed to assess the impact of this mutation across populations considering that already some rivers have about 10% of “apparent” XY sex reversed fish (Cavileer et al., 2015; Williamson et al., 2008). Interestingly, Cavileer et al published a Chinook salmon *sdY* genomic DNA sequence (GenBank: KC756279) from Tozitna River, Alaska and a *sdY* cDNA sequence (GenBank: KF006343) from embryonic males from Clearwater River, Idaho (Cavileer et al., 2015). The alignment of those sequences with ours revealed the same nucleotide substitution (A/T) in exon 3 but not the (A/G) substitution in exon 2. We also report in the present study RNA-Seq data from a Chinook male from the Umatilla river in Oregon that also has these two mutations. The presence of the deleterious mutation in different populations across the North America coast (Alaska, Oregon and California) support the hypothesis that this mutational event occurred before the establishment of these different populations, and that this mutation could be widespread in many Chinook populations. The inactive *sdY* copy as a non-functional gene should accumulate further mutations quite rapidly and should show many features of gene decay. However, the propagation of the mutation requires that sex reversed females maintain similar reproductive capacity and fitness as wild type (Senior *et al*. 2012) or that the Y-chromosome accumulates more copies by genetic drift. Another point of interest is the origin of Y-chromosome which has only the defective *sdY* version. The Chinook Y+ has two copies of sdY, wildtype and the defective sdY version. The salmonid sex determination locus has been assigned features of a « jumping locus » behaving like a giant mobile element (Faber-Hammond *et al*. 2015). Thus, a local gene duplication is a very likely origin of both copies creating the Y+. In a second event, one of the *sdY* copies acquired the inactivating mutation and the silent mutation. The deletion of the functional *sdY* from the Y+ resulted in the formation of the Y-. The loss of the male determining function from the Y- makes it segregate like an X. It is tempting to speculate that the defective *sdY* will disappear from the Y- one day and similarly from the Y+. The occurrence of one male in our crosses that was genotyped to contain only the wildtype version of *sdY* could support such a scenario. The question may arise if the hypothetical ancestral Y+ still exists in the wild Chinook populations. The implementation of an accurate assay such as PCR or qPCR followed by sequencing to test the presence/absence of the mutation will be helpful for both aquaculture, stock management, population genetics and conservation biology of this species (Yano *et al*. 2013).

In conclusion, we demonstrated that some wild Chinook salmon harbor a copy of the sex determining gene sdY gene, which, due to a missense mutation, lost the ability for testis determination and explains the genetic status of XY “apparent” male to female sex reversals. We show this mutant Y is not effective anymore and behaves like an X-chromosome. So far except for *sdY*, no gene promoting maleness expressed as early as *sdY*, has been identified in the Y-specific region of salmonids. Also, no gene present on the X has been shown to be lost from the male-specific on the Y chromosome (MSY). On this basis the XY females are not bona-fide sex reversals. Their genome does not harbor a gene that would induce male development, thus their phenotype reflects accurately their genotype.

## MATERIALS AND METHODS

### Chinook samples genotyping

Family panels and genetic samples were the same as the ones described in Williamson and May (Williamson and May 2005), and were produced from a fall-run Californian Chinook population harvested at the Merced Hatchery. PCR sequencing analysis of exon 2, intron 2 and exon 3 was performed using a long-range PCR protocol and primers designed upstream and downstream of the *sdY* Chinook gene sequence (GenBank ID = KC756279.2), followed by targeted Sanger resequencing of these PCR fragments with internal primers for exon 2, intron 2 and exon 3. Long-range PCR were carried out in a final volume of 50 μl containing 0.4 μM of each primers (sdYChinook-F2: TTGGCTCCCAGGAAAACATTTCT; sdYChinook-R1: CAGAACAAACAGCATGAAGTAAGCA), 80 ng gDNA, 1X of 10X AccuPrime^™^ buffer II (including dNTPs), and 1.5 μl per reaction of AccuPrime^™^ HiFi Taq DNA polymerase. Cycling conditions were as follows: 94 °C for 1 min, then 35 cycles of (94 °C for 30 sec + 64 °C for 30 sec + 68 °C for 6 min).

### Chinook testis RNA-seq

Chinook testis was sampled from an adult male from the Umatilla river (OR), and the testis library was prepared using the TruSeq RNA sample preparation kit, according to manufacturer instructions (Illumina, San Diego, CA) as previously described (Pasquier et al. 2016). These testicular transcriptome reads were mapped on a female Chinook genome assembly (Otsh_v1.0, GCA_002872995.1) plus the *sdY* Chinook gene sequence (GenBank ID = KC756279.2) using BWA (Li and Durbin 2009) with stringent mapping parameters (maximum number of mismatches allowed –aln 2). High quality reads (MAPQ > 40) remapping on the *sdY* gene were visualized and analyzed with the IGV software (Robinson *et al*. 2011).

### Protein structure prediction

The 3D model of SdY-I183N was predicted using the X-ray structure of the dimeric interferon regulatory factor 5 transactivation domain at 2 Å resolution (PDB ID 3DSH) as template (Chen et al., 2008). The three-dimensional views of SdY-N183 were obtained with PyMOL software (Molecular Graphics System, Version 1.7.4 Schrödinger, LLC).

### Cloning

Plasmids and primers used are listed in supplementary Table 2–3. A forward primer was generated from the coding sequence of the rainbow trout SdY with a point mutation T/A to mimic the SdY-I183N mutation. Next, the amplified fragment containing the mutation was inserted in pCS2^+^-FLAG:SdY. From this plasmid, a PCR-amplified fragment corresponding to SdY-N183 was inserted into pCS2^+^, pCS2^+^-3xHA, pCS2^+^-3xFLAG, pGEX-4T1 expression vectors. The pCS2^+^-3xHA:emGFP:SdY-N183 plasmid was obtained by inserting a PCR-amplified fragment corresponding to emGFP in-frame into the EcoRI site between 3xHA and SdY-N183.

### Cell culture

Human embryonic kidney (HEK 293T) cells were cultured and maintained in DMEM medium (PAN Biotech), supplemented with 10% FCS (PAN Biotech) and 1% Penicillin-Streptomycin (PAN Biotech) at 37°C with 5% CO_2_. HEK 293 transfections were performed by incubating cells with Polyethylenimine (PEI) (100 mg/mL PEI diluted 1:100 in 150mM NaCl) and the respective plasmids (10 μg for 10 cm dishes, 2 μg for 6-well plates) for 6-8 hours into fresh medium. Then, the medium was discarded and fresh medium was added.

### Immunofluorescence

HEK 293T cells were seeded on 6-well plates containing coverslips. After transfection of the corresponding plasmids (pCS2^+^-SdY-N183; pCS2^+^-FLAG:SdY-N183; pCS2^+^-3xFLAG:SdY-N183; pCS2^+^-HistoneH2B:mCherry)) with or without (pCS2^+^-HA-mCherry-Foxl2b2) for 48 h, cells were fixed in 4% fresh paraformaldehyde (PFA) for 15 min, extensively washed, and permeabilized with 0.1% Triton X-100 in PBS for 10 min. Then cells were blocked with 1% BSA during 20 min. Primary antibody (Supp Table 4) was incubated overnight at 4°C. After extensive washes with PBS, cells were incubated with Alexa 488 conjugated secondary antibodies in 1% BSA for 1 h, followed by Hoechst 33342 (Invitrogen) staining for 5 min (1 μg/mL final concentration). Cells were mounted using Mowiol 4-88 (Roth). Confocal images were acquired with a Nikon Eclipse C1 laser-scanning microscope (Nikon), fitted with a 60x Nikon objective (PL APO, 1.4 NA), and Nikon image software. Images were collected at 1024×1024 pixel resolution. The stained cells were optically sectioned in the z axis. The step size in the z axis varied from 0.2 to 0.25 mm to obtain 50 slices per imaged file. All experiments were independently repeated several times at least three times. Cytoplasmic localization was counted when the main source of signal comes from the cytoplasm. A nucleocytoplasmic localization was counted when a strong signal was detected in both cytoplasm and nucleus. In a same way, a nuclear localization was counted when the signal was detected in the nucleus and when the signal follows the pattern of fluorescence intensity.

### Western Blotting

Cells were lysed in a HEPES-based lysis buffer (20 mM HEPES (pH 7.8), 500 mM NaCl, 5 mM MgCl2, 5 mM KCl, 0.1% deoxycholate, 0.5% Nonidet-P40, 10 mg/ml aprotinin, 10 mg/ml leupeptin, 200 mM sodium orthovanadate, 1 mM phenylmethanesulphonylfluoride and 100 mM NaF) for 3 h. Cell debris was pelleted by centrifugation for 15 min at 16000 g. Cell lysate protein concentration was measured with a Bradford assay (Cary 50 Spectrophotometer, Varian). The protein lysates (30–50 μg) were resolved by SDS-PAGE on 12% Tris-glycine gels followed by transfer to nitrocellulose membranes. Unspecific binding was blocked with 5% BSA in TBST (10 mM Tris pH 7.9; 150 mM NaCl; 0.1% Tween) for 1h at room temperature. Incubation with primary antibodies was performed overnight at 4°C. After three washes with TBST, HRP conjugated antibodies were incubated with blocking solution for 1h. Following the washes, membranes were incubated with SuperSignal West Pico Chemiluminescent Substrate (Thermo Scientific) for 1 min. The signal from the membranes was detected using the Photo Image Station 4000MM (Kodak). At least two independent experiments were performed and representative protein blot images are shown. Quantitative analysis was performed with ImageJ 1.48v software (www.imagej.nih.gov).

### Cycloheximide treatment

HEK 293T cells were transfected either with 3xHA-SdY or 3xHA-SdY-N183 with or without the 3xFLAG-tFoxl2b2 expression vector. 48 h post-transfection, cells were treated with 50 μM of the protein synthesis inhibitor cycloheximide (Calbiochem), or ethanol as vehicle control during 4 h or 8 h. Untreated cells (0 hour) and treated cells were harvested and subjected to cell lysis followed by SDS-PAGE and Western Blot as described above.

### MG132 treatment

HEK 293T cells were transfected either with 3xHA:SdY or with 3xHA:SdY-N183 with or without the 3xFLAG:Foxl2b2 expression vector. 48 hours post-transfection, cells were treated with 20 μM of a proteasome inhibitor, i.e., MG132 (Merck) or DMSO as vehicle control during 8 h. Untreated cells (0 hours) and treated cells were harvested and subjected to cell lysis followed by SDS-PAGE and Western blot as described above.

### Luciferase assay

HEK 293T cells were transfected using PEI with the following plasmids: 0.3 μg of pGL3-*Olacyp19a1a* sequence (kindly provided by D. Wang Deshou); 0.05 μg-0.4 μg of pCS2^+^-SdY-N183 expression plasmid; 0.05-0.4 μg of pCS2^+^-OlaFoxl2; 0.1 μg of pcDNA3.1-OlaNr5a1 and 0,1 μg of pTK-Renilla used for calibration. Each experiment was performed with 1.0 μg final amount. Adjustments were made with empty vector (pCS2+) accordingly. Firefly luciferase and *Renilla* luciferase readings were obtained using the Dual-Luciferase Reporter Assay System (Promega) and LUMAT LB 9501 luminometer (Berthold Technologies GmbH & Co. KG, Bad Wildbad, Germany).

### Statistical analysis

Data were analyzed using a two-sided unpaired Student’s t-test. Additionally, luciferase assay was subjected to one-way ANOVA with post-hoc Dunnett tests. Significant differences are symbolized in figures by asterisks if p<0.001 (***), p<0.05 (**), p<0.01 (*) or N.S. if not significant.

## Author Contributions

S. B., A.H., E.J., A.Y. performed experiments and analyzed the data, J. B., H. P., L. J., R. G., K.W, M.M, P.S. provided samples, T. M. modelled the SdY and IRF proteins, Y. G and M.S. designed the study and supervised the project, S.B., A.H., G. M., Y.G., and M.S. wrote the paper.

## Acknowledgements

We acknowledge Wang Deshou for the *olacyp19a1a* promoter. This work was supported by Agence Nationale de la Recherche (ANR) grant ANR ANR-11-BSV7-0016 (SDS project) and grants to MS by the Deutsche Forschungsgemeinschaft (Scha408/12-1, 10-1).

## Data availability

The Chinook testis RNA-Seq sequences are available at GenBank Sequence Read Archive (SRA) under the accession number: SRX4998097. Supplementary Table 1 is available via the GSA Figshare portal.

## Supplemental Information

**Supplemental Figure 1.**
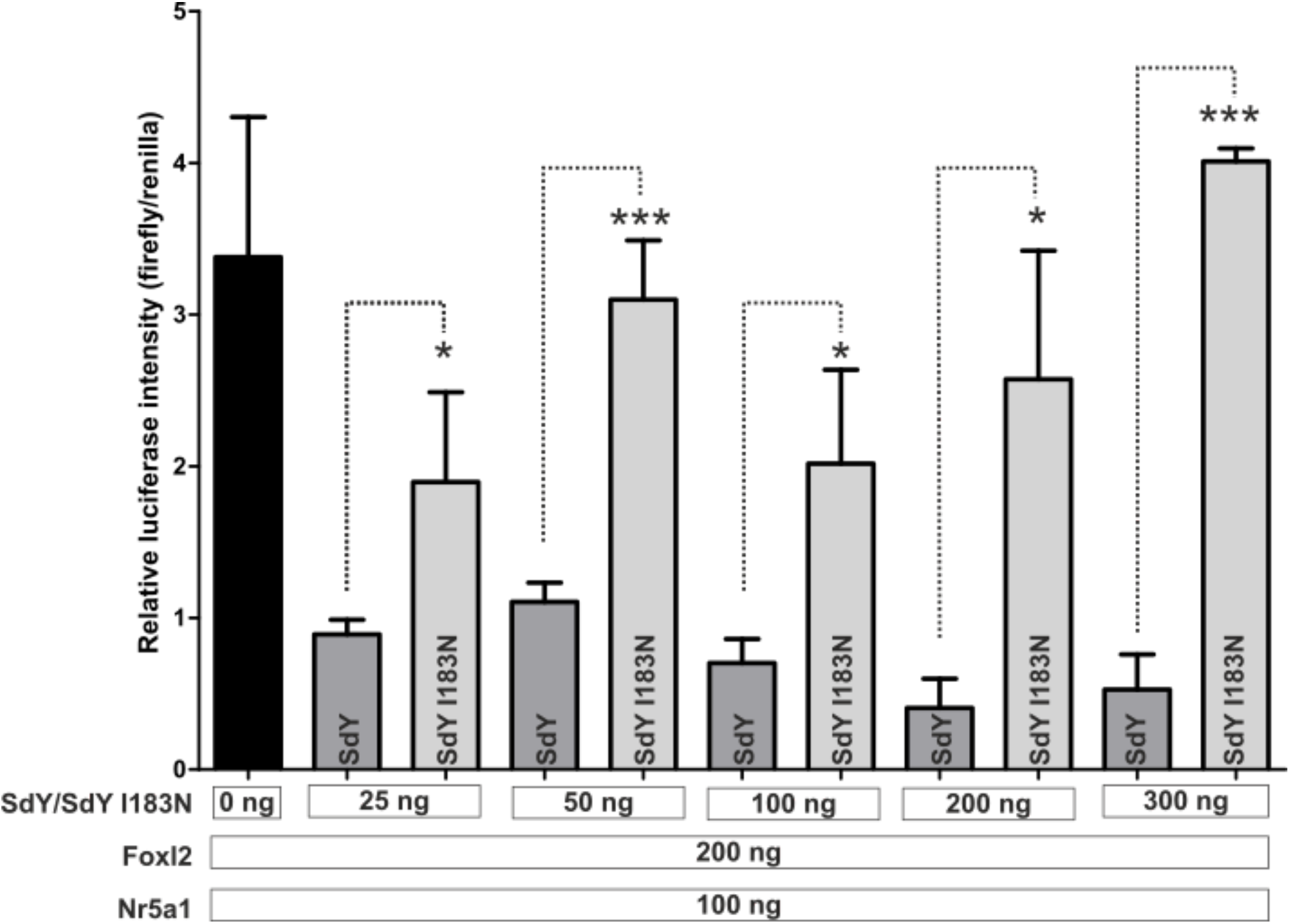
Comparison of SdY wild type and SdY I183N effect on Nr5a1/Foxl2 induced *cyp19a1a* promoter. Data obtained in Figure 5 were compared with data from (Bertho *et al*. 2018). Statistical significances of activity changes between Foxl2 (200 ng) / Nr5a1 (100 ng)/ SdY (25 to 300 ng) and Foxl2 (200 ng) / Nr5a1 (100 ng)/ SdY I183N (25 to 300 ng) (two-sided,Student’s *t*-test) are shown by asterisks, p<0.05 (*); p<0.001 (***).

**Supplementary Table 1.** Genotype/phenotype ratios based on PCR assay of exon 2, intron 2 and exon 3 for *sdY* amplification in male and female from three crosses normal family 84 x B; XY family 126xD; XY family 118xC. (available via the GSA Figshare portal).

**Supplementary Table 2.** Plasmids

| Plasmid | Description |
| --- | --- |
| **pCS2+** | Empty expression vector under a CMV promoter |
| **pCS2<sup>+</sup>-3xHA** | Cloning of the 3xHA tag sequence in pCS2<sup>+</sup> sequence |
| **pCS2<sup>+</sup>-3xFlag** | Cloning of the 3xFlag tag sequence in pCS2<sup>+</sup> sequence |
| **pCS2<sup>+</sup>-HA:mCherry** | Cloning of a HA:mCherry tag in pCS2<sup>+</sup> plasmid (gift from M. Gessler (University of Wuerzburg)) |
| **pCS2<sup>+</sup>-SdY-N183** | SdY-N183 CDS cloned into a pCS2<sup>+</sup> backbone from pFLAG:SdY:I183N plasmid |
| **pCS2<sup>+</sup>-FLAG:SdY-N183** | FLAG:SdY fusion under the control of a CMV promoter, SdY-N183 CDS cloned into the pCS2<sup>+</sup> backbone. |
| **pCS2<sup>+</sup>-3xFlag :SdY-N183** | 3xFLAG:SdY-N183 fusion under the control of a CMV promoter, SdY-N183 CDS cloned into the pCS2<sup>+</sup>-3xFLAG backbone. |
| **pCS2<sup>+</sup>-HA :mCherry:Foxl2b2** | HA:mCherry:Foxl2b2 fusion under the control of a CMV promoter, Foxl2b CDS cloned into a pCS2<sup>+</sup>-HA:mCherry backbone. |
| **pCS2<sup>+</sup>-Ol-Foxl2** | Cloning of Oryzias latipes Foxl2 CDS from pCS2<sup>+</sup>-OlaFoxl2:FLAG |
| **pcDNA3.1-OlaNr5a1** | Gene synthesis of medaka Nr5a1 by between the EcoR1-Xho1 restriction enzyme site ( genescript) |
| **pGL3 basic** | Vector backbone containing a cDNA modified firefly luciferase (luc+) |
| **pGL3-Ol-cyp19a1a** | Firefly luciferase under the control of two kb upstream the transcription start site of Ol-Cyp19a1a |
| **pminiTK-Renilla** | Renilla luciferase under the control of a mini-thymidine Kinase promoter |

**Supplementary Table 3.**
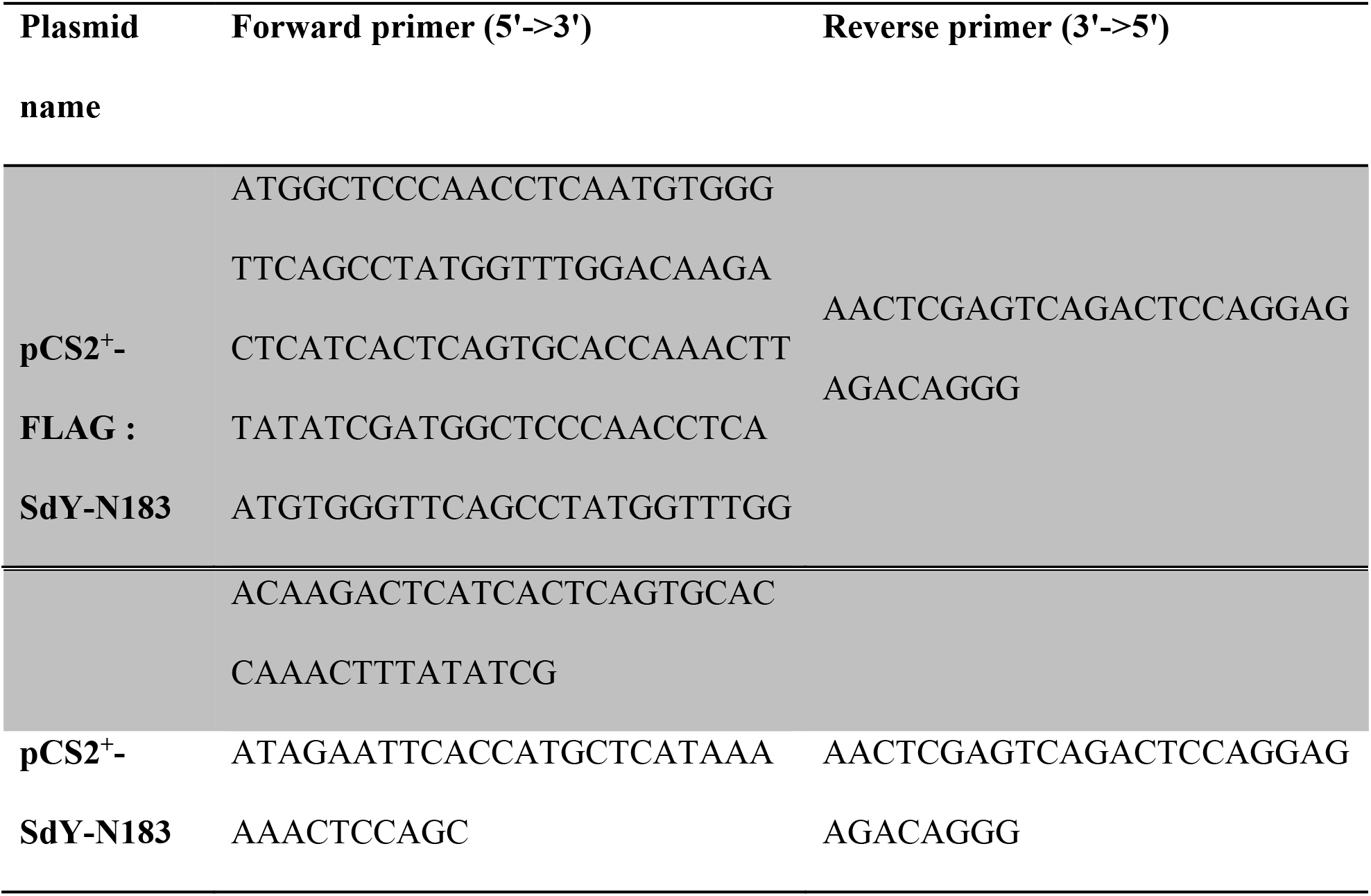
Primers

**Supplementary Table 4.**
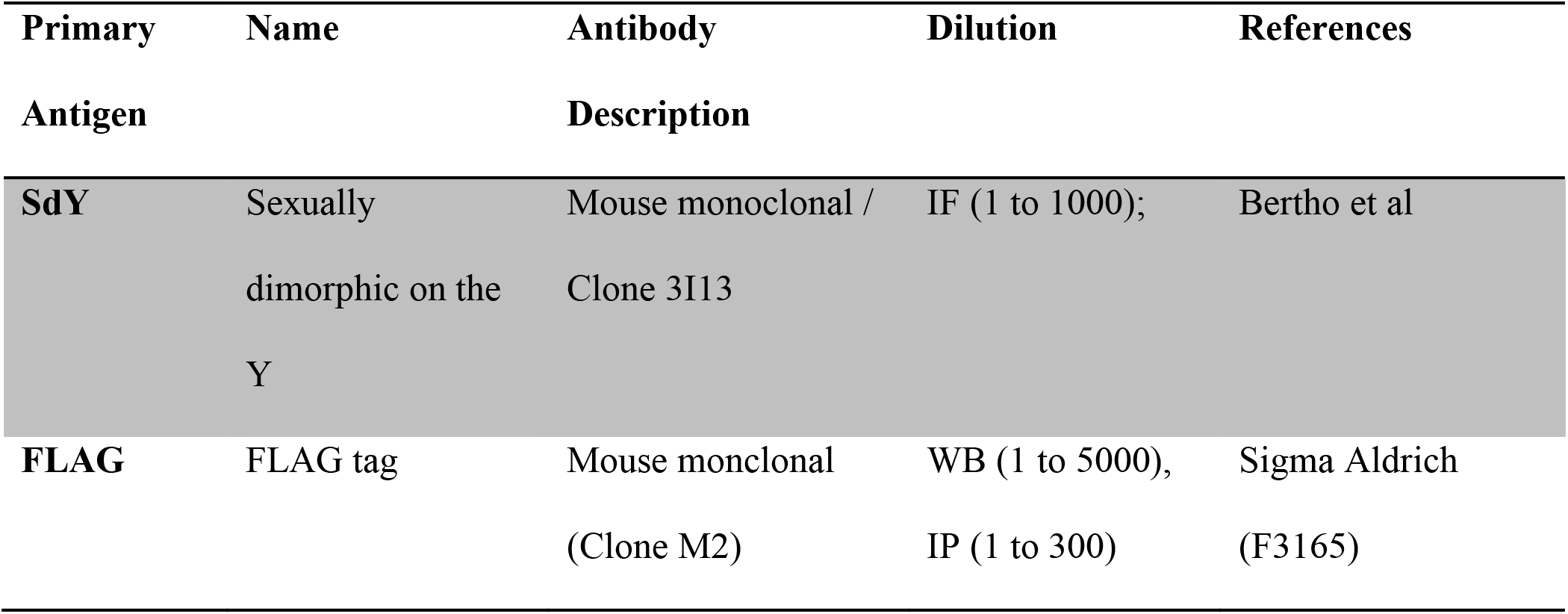
Antibodies used in this study.

